# Generalized Euler-Lotka equation for correlated cell divisions

**DOI:** 10.1101/2021.01.11.426278

**Authors:** Simone Pigolotti

## Abstract

Cell division times in microbial populations display significant fluctuations. These fluctuations impact the population growth rate in a non-trivial way. If fluctuations are uncorrelated among different cells, the population growth rate is predicted by the Euler-Lotka equation, which is a classic result in mathematical biology. However, cell division times can present significant correlations, due to physical properties of cells that are passed from mothers to daughters. In this paper, we derive an equation remarkably similar to the Euler-Lotka equation which is valid in the presence of correlations. Our exact result is based on large deviation theory and does not require particularly strong assumptions on the underlying dynamics. We apply our theory to a phenomenological model of bacterial cell division. We find that the discrepancy between the growth rate predicted by the Euler-Lotka equation and our generalized version is relatively small, but large enough to be measurable in experiments.

---

Microbial populations in steady, nutrient-rich conditions tend to grow exponentially. Their exponential growth rate Λ can be taken as a proxy for the population fitness and is therefore a biologically important quantity. In a population of cells dividing at regular times *τ*, the population size at a time *T* multiple of *τ* is *N*(*T*) = *N*(0)2^*T/τ*^, so that Λ = ln 2/*τ*. In practice, cell division times of microbial populations significantly fluctuate, so that the division time *τ_i_* of a given individ-ual *i* must be considered as a random quantity. As a consequence, the growth of *N*(*T*) is stochastic. In these cases, we can still define an exponential growth rate by

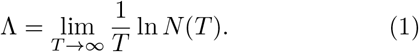

For independent, identically distributed cell division times, the exponential growth rate converges to a deterministic value and can be computed as solution of the celebrated Euler-Lotka equation

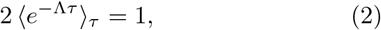

where we denote the average over the distribution *p*(*τ*) of the division times by 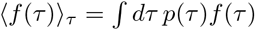. We use this notation also for discrete variables, with the integral appropriately replaced by a sum. Equation (2) is a classic result in mathematical biology. A recent experimental study has tested its prediction by tracking individual cell divisions in a microfluidic device [1]. Beside microbial populations, the Euler-Lotka equation finds important applications in epidemiology, where the factor 2 is replaced by the reproductive number *R*_0_ [2].

Experimental studies have revealed that fluctuations in microbial cell features are correlated among generations [3–5]. These correlations are caused by properties of cells that are passed through generations. These properties can be physical such as cell mass, or biological such as gene expression. Their fluctuations are often controlled to preserve homeostasis, i.e., a stable state of cells across generations. For example, experimental and theoretical studies provided evidences for an “adder” mechanism, in which cells attempt at growing their mass by a constant amount before dividing [4, 6, 7].

Regardless of the mechanism underlying correlations in cell division times, generalizing Eq. (2) to the correlated case has proven to be a hard problem. One relatively simple case is the “Markovian” scenario where a cell generation time conditionally depends only on that of her mother. Expressions for the growth rate in these cases have been derived in classic works by Powell [8] and Lebowitz and Rubinow [9], see also [10]. Alternative approaches estimate the growth rate by comparing the outcome of sampling the population forward in time with retrospective sampling, in which individuals in the final population are traced back to their ancestors [11, 12]. A recent study linked the exponential growth rate Λ to the asymptotic distribution of the number of cell divisions Δ among lineages [13]. This approach makes use of techniques borrowed from large deviation theory and has the advantage of neither requiring the Markovian assumption, nor retrospective sampling.

In this paper, we introduce a generalization of the Euler-Lotka equation (2) which is valid for correlated cell division times:

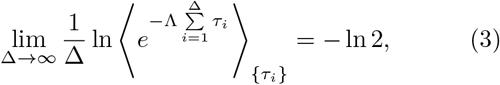

where 〈…〉{_*τ_i_*}_ denotes an average over sequences {*τ_i_*} of cell division times along lineages. If the *τ_i_*s are uncorrelated, then the left hand side of Eq. (3) reduces to the cumulant generating function ln〈*e^qτ^*〉_*τ*_, and therefore Eq. (3) becomes equivalent to the traditional Euler-Lotka equation (2). Equation (3) only requires as hypotheses that population dynamics is at steady state and the sum of the *τ_i_*s across a lineage satisfies a large deviation principle with a convex rate function, which are in practice rather mild assumptions. Equation (3) can therefore be used to compute the population growth rate from individual cell division times in rather general settings.

We consider a microbial population initially constituted of a single individual. The population grows in time by a sequence of cell divisions. We represent the genealogy of the population by a tree, whose nodes are cell division events and branches are times between consecutive cell divisions, see Fig. 1a.

**FIG. 1.**
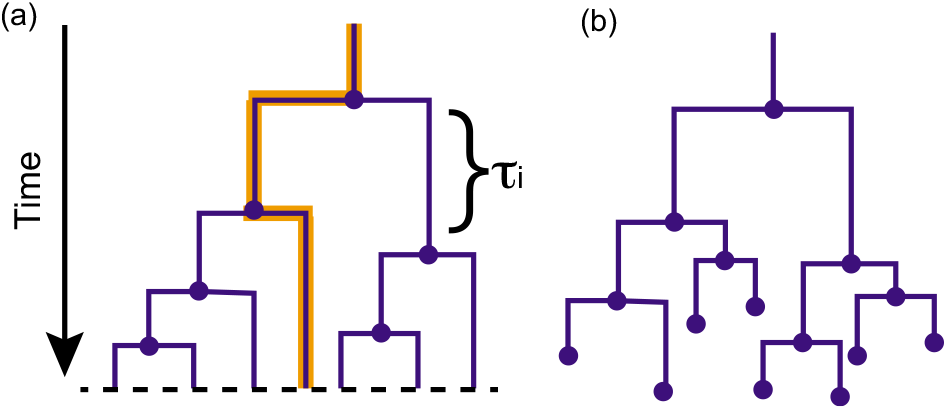
Population dynamics represented as a lineage tree. (a) a microbial population grows in time from a single cell. Nodes (circle) denote cell division events. Lengths of branches denote the cell division times *τ_i_*s. One lineage is represented with a thick orange line. In this case, the population is let to evolve until a fixed time *T*. (b) Lineage tree in an alternative ensemble, in which lineages are let to evolve until they have accumulated exactly Δ = 4 cell divisions.

We now formalize the concept of a lineage. A lineage is identified by an individual in the population at time *T* complemented by its past history, i.e., the number Δ of cell divisions separating it from the individual at time 0 and the sequence of cell division times {*τ_i_*} of all its ancestors, see Fig. 1a. Following Refs. [13, 14], we now imagine to randomly select a lineage by starting from the initial individual and picking at each divisions one of the two newborns with equal probability. With this procedure, a lineage that underwent Δ cell divisions is chosen with probability 2^−Δ^. The probability that a randomly selected lineage includes Δ cell division events is equal to *p*(Δ;*T*) = 2^−Δ^*N*(Δ; *T*), where *N*(Δ; *T*) is the number of lineages with Δ cell divisions at time *T*. Since 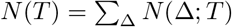, we obtain

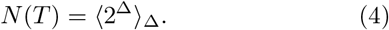

Substituting this expression into the definition of the exponential growth rate, Eq. (1), we find

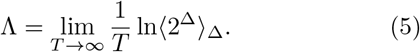

We remark that, in this argument, we did not distinguish between the a priori probabilities *p*(Δ; *T*) and the empirical ones, estimated from lineage frequencies in a single tree. In fact, it can be shown that the empirical probabilities rapidly converge to the a priori ones in the large *T* limit [13, 15], so that this distinction is not important for our aims.

To make further progress, we introduce some ideas from large deviation theory [16]. Large deviation theory describes the leading behavior of distributions when a parameter (like the time *T* in our case) becomes large. In large deviation theory, variables such as Δ, whose average is proportional to *T*, are called extensive. We associate with *T* the intensive variable *δ* = Δ/*T*, whose average tends to a constant for large *T*. The large deviation principle for *δ* is expressed by

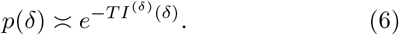

The function *I*^(*δ*)^ (*δ*) is called the rate function. We use the notation *I*^(*δ*)^ to stress that I is the rate function associated with the distribution of the variable *δ*. The symbol 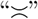 denotes the leading exponential behavior; it can be seen as a shorthand for *I*(*δ*) = – lim_*T*→∞_[ln *p*(*δ*)]/*T*.

An alternative way of studying asymptotic fluctuations of intensive random variables is via the scaled cumulant generating function, defined by

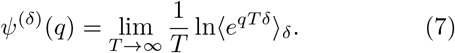

The Gartner-Ellis theorem states that, if the rate function is convex, it is related with the scaled cumulant generating by a Legendre-Fenchel transform

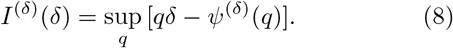

Since the Legendre-Fenchel transform is an involution, it also holds that *ψ*^(*δ*)^(*q*) = sup_*δ*_ [*qδ* – *I*^(*δ*)^(*δ*)].

We now return to Eq. (4) and briefly summarize the main result of Ref. [13]. Assuming that *δ* satisfies a large deviation principle, we obtain

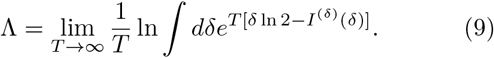

In the limit *T* → ∞, the integral can be evaluated with the method of steepest descent, obtaining

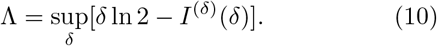

Equation (10) is the central result of Ref. [13]. An alternative way to obtain it is to directly identify the expression of the scaled cumulant generating function in Eq. (5):

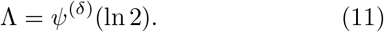

Equation (10) then follows by expressing the scaled cumulant generating function in terms of the rate function by means of the Gartner-Ellis theorem.

Application of this theory requires knowledge of the asymptotic distribution of Δ, or its intensive counterpart δ. However, in analogy with the Euler-Lotka equation (2) it would be desirable to express the growth rate in terms of the distribution of division times and its correlations. To this aim, we now consider a case in which, rather than letting the population grow until a given time *T*, each lineage is let to grow until it has accumulated exactly Δ cell divisions, see Fig. 1b. In this alternative *ensemble*, Δ is fixed whereas *T* fluctuates among lineages. In this case, we consider *T* as an extensive random variable, since its average grows linearly with the fixed large parameter Δ. We similarly associate with *T* the intensive variable *t* = *T*/Δ. We expect *t* to satisfy as well a large deviation principle:

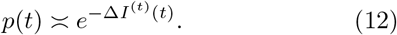

In the language of probability theory and in particular of queuing theory, Δ(*T*) is called a counting process and *T*(Δ) its inverse. A useful result [17] states that the large deviation of their associated intensive variables, *δ* and *t* respectively, are related by

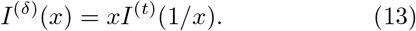

Note that this is the result that one would obtain by taking the large deviation form of the probability distribution and simply applying the rules for a change of variable.

We now substitute this result into Eq. (10), obtaining

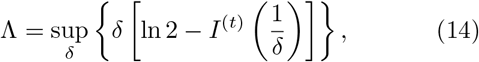

and, by applying the Gartner-Ellis theorem,

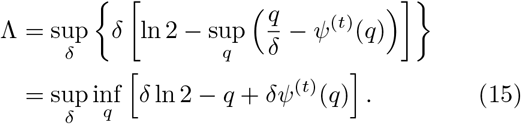

We assume that the function in curly brackets smoothly depends on *δ* and *q* and therefore compute the supremum and infimum by simply taking derivatives. The extremality condition respect to *δ* is expressed by

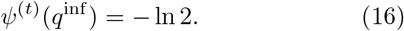

Substituting this condition back into Eq. (15) yields Λ = –*q*^inf^, so that we rewrite Eq. (16) as

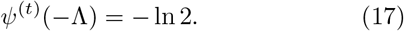

Upon substituting the definition of the scaled cumulant generating function, Eq. (7), into Eq. (17), we obtain the generalized Euler-Lotka equation (3), as anticipated.

Taking the derivative in Eq. (15) respect to *q* results in

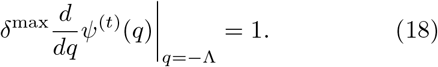

This equation relates the dominant value of *δ* with the statistics of the division times and provides another facet to the generalized Euler-Lotka theory. Equation (18) is best interpreted in the simple case of uncorrelated cell divisions, where it reduces to *δ*^max^ = 1/(*τe*^−Λ*τ*^). If Λ ≪ 1, the dominant value of γ is simply its average value, i.e. the inverse of the average division time. However, for quickly growing population, the dominant value of δ becomes significantly larger than this value, as cells that reproduce faster contribute more to population growth.

To illustrate this result, we consider a phenomenological model of bacterial growth inspired to the “adder” principle [4]. In the model, each bacterial cell grows in length at a rate *α*. The rate *α* fluctuates among cells but is constant in time per each individual cell. is distributed according to the formula

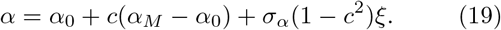

where *α_M_* is the value of *α* of the mother of the considered cell. The parameters *α*_0_ and *σ_α_* are the average and variance of the distribution of α, respectively. The parameter *ξ* is a Gaussian random variable with zero average and unit variance. Finally, the parameter *c* controls the degree of correlations between the growth rate of mothers and daughters.

Each cell is characterized by a length *s_b_* at birth and *s_d_* at death. The adder model postulates that the added length *l* = *s_d_* – *s_b_* is roughly constant among cells. After division, a daughter inherits a fraction *f* of the mother’s length. We allow for some variability by taking both *f* and *l* as random variables, with Gaussian and lognormal distribution respectively. Averages and variances of these distributions are estimated from experimental data [4]. According to our assumptions, the time between cell divisions is expressed by *τ* = [*α* ln(*s_d_*/*s_b_*)]^-1^.

We simulate the model to obtained *n*_lin_ lineages, each including Δ cell divisions. We plug the results of these simulations into the generalized Euler-Lotka equation (3) and thereby compute the value of Λ, see Fig. 2. We consider two different scenarios: one in which the growth rate *α* is positively correlated across generations *c* = 0.5 and one in which it is uncorrelated (*c* = 0). In both scenarios, we compare the results of the generalized Euler-Lotka equations with those of the classic Euler-Lotka equation (2). We find that, in the correlated case, the growth rates predicted by the two equations are nearly indistinguishable, see Fig. 2a and 2b. This is not surprising, as the adder mechanism tends to cause negative correlations among cell division times, which can be counterbalance by positive correlations in the growth rate [4]. Indeed, when α is uncorrelated, the adder mechanism is not compensated and we find a difference between the growth rate predicted by the two equations, see Fig. 2c and 2d. In this case, this difference is on the order of 0.5%. Our results show that such a small difference can be detected by our framework by tracking, for example, a few hundred lineages over a time corresponding to about 20 cell divisions. This implies that our theory is practically applicable to modern experimental data.

**FIG. 2.**
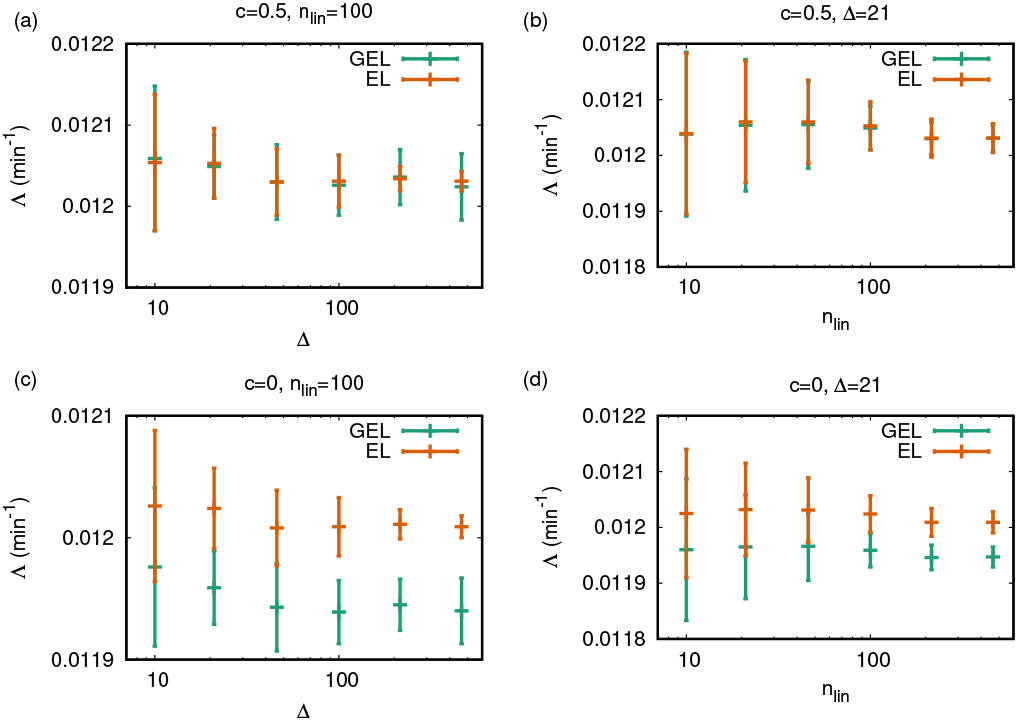
Estimates of the growth rate Λ in the adder model of Ref. [4] obtained by the generalized Euler-Lotka equation (3) (GEL) and by the conventional Euler-Lotka equation (2) (EL). In all panels, parameters of the growth rate distribution are *α*_0_ = 0.0255 min^−1^ and *σ_α_* = 0.0027 min^−1^. The inherited length fraction *f* is distributed according to a Gaussian with mean *f*_0_ = 0.5 and standard deviation *σ_f_* = 0.03. The added length *l* follows a lognormal distribution with mean *l*_0_ = 3.21 *μm* and standard deviation *σ_l_* = 0.54 *μm*. In panels (a) and (c), we fix the number of lineages *n*_lin_ = 100 and plot the results as a function of the number of cell divisions Δ in each lineage. In panels (b) and (d), Δ = 21 is fixed and results are plotted as a function of *n*_lin_. Panels (a) and (b): growth rates of mothers and daughters are positively correlated (*c* = 0.5). Panels (c) and (d): growth rates of mothers and daughters are uncorrelated (*c* = 0). In all panels, simulations are repeated 20 times; error bars denote standard deviations computed from these realizations.

An advantage of this approach is that it allows to use the arsenal of techniques from large deviation theory [16] to compute the scaled cumulant generating function, and thereby the growth rate via Eq. (3). For example, we apply our theory to the Markovian case in which the division time conditionally depends on the maternal division time only, i.e.,

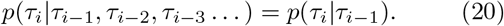

In this case, one has

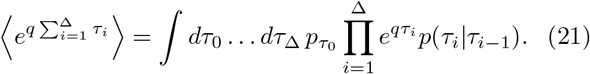

This expression and the definition of the scaled cumulant generating function, Eq. (7), imply that

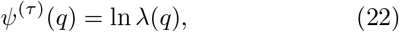

where λ(*q*) is the leading eigenvalue of the convolution kernel

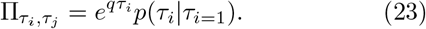

Combining Eq. (3) and Eq. (22) we obtain 2λ(–Λ) = 1, which is a classic result for the Markovian case [8–10].

In conclusion, in this paper we derived the generalized Euler-Lotka equation (3). This equation describes the growth rate of populations where cell divisions occur in a correlated way. We obtained this result by means of a result in queuing theory [17] that was recently applied in stochastic thermodynamics [18] and to study enzyme replicating information [19]. Using a phenomenological model of bacterial cell division, we have demonstrated that our result can be easily applied to lineage data. A comparison with the prediction of the traditional Euler-Lotka equation permits to quantitatively assess the impact of correlations on the population growth rate. Due to these properties, we expect the generalized Euler-Lotka equation to become a useful tool to analyze population dynamics tracked at the single-cell level in experiments.

I thank Deepak Bhat and Anzhelika Koldaeva for discussions on the Euler-Lotka theory and Luca Peliti for a critical reading of the manuscript.

